# Novel Genes and Polymorphisms in Human Immunoglobulin Light Chains Across Diverse Populations Through Comprehensive IMGT Analysis

**DOI:** 10.1101/2025.09.25.678462

**Authors:** Maria Georga, Ariadni Papadaki, Géraldine Folch, Joumana Jabado-Michaloud, Guilhem Zeitoun, Patrice Duroux, Véronique Giudicelli, Sofia Kossida

## Abstract

The human immunoglobulin light chain loci, kappa (IGK) and lambda (IGL), are structurally complex genomic regions with germline gene content that is not yet fully characterized. These loci are marked by extensive gene duplication, allelic diversity, and segmental duplications, features that contribute critically to the adaptive immune response. In this study, we present a comprehensive IMGT annotation of IGK and IGL using two high-quality human reference assemblies (GRCh38 and T2T-CHM13) along with 142 and 125 additional chromosomal-level haploid assemblies, respectively for each locus, from individuals representing all major human superpopulations. Detailed gene and allele annotation of the reference assemblies led to the identification of 5 novel IGKV genes and 8 new IGKV alleles, 16 new IGLV genes, and 22 novel IGLV alleles. These were confirmed through assembly read validation, presence in whole genome sequencing datasets, and recurrence in multiple assemblies. Gene-level identification across the broader dataset enabled assessment of structural variation (SV) at both loci. IGL displayed high conservation, with recurrent absence observed for only one gene. In contrast, IGK exhibited greater variability, including complete loss of the distal region in certain assemblies. This structural diversity was analyzed across superpopulations, allowing us to map potential patterns of gene presence and absence across different ancestral groups. All newly identified genes were consistently observed across individuals and genomic backgrounds. This work enhances the structural resolution of the IGK and IGL loci and expands the IMGT reference directory with newly described germline genes and alleles. The results provide a more complete view of light chain genomic diversity and serve as a valuable resource for studies of antibody gene repertoires, immunogenetic variation, monoclonal antibody development and population-level diversity.

## Introduction

The human immunoglobulin light chains, encoded by the kappa (IGK) and lambda (IGL) loci, are essential for antibody function and a robust adaptive immune response (1). Along with the heavy chain (IGH), one of the light chain loci will form the antigen-binding site of immunoglobulins (IG), influencing their specificity, affinity, and molecular characteristics. Combinatorial pairing of heavy and light chains significantly expands the diversity of the human antibody repertoire. Yet, despite their central role in immunity, the genomic architecture and germline content of the IGK and IGL loci remain incompletely resolved, particularly regarding structural variation and diversity across individuals from diverse populations.

In contrast to the single, contiguous heavy chain locus (IGH) locus, the light chain genes are distributed across two distinct chromosomal regions: IGK on 2p11.2 and IGL on 22q11.2 (Supplementary Figure S1A and S1B) (2). These two loci contain multiple variable (V), joining (J), and constant (C) genes and are organized to support the generation of IG diversity through somatic VJ recombination, where a single V gene is joined to a J gene and then the V-J segments is spliced to a C for a functional light chain. The IGK locus exhibits a unique organization, consisting of two large, inverted, highly similar IGKV clusters containing V genes, followed by the J-C genes. The cluster closer to the J-C genes is termed proximal, while the more distant one is distal. In contrast, IGL features a tandem array of IGLV genes followed by J-C clusters (3). Although the two loci differ in genomic organization, both are characterized by high sequence similarity among their V genes and among their J genes, and both loci display extensive allelic diversity. These characteristics underpin their complex structure and present substantial challenges for accurate genomic assembly and annotation.

Despite their essential role in antigen recognition and their extensive use in antibody-based therapeutics, the structural complexities of the light chain loci have resulted in significant gaps in our understanding of germline gene content (4). Few studies have aimed to uncover new polymorphisms, copy number variations, and structural variants in IGK and IGL, and even fewer have done so on a large scale across individuals from diverse backgrounds to assess genetic diversity and potential patterns.

Reference genomes such as GRCh38 (5) capture only a portion of IGK (chromosome 2) and IGL (chromosome 22) loci, and the high similarity between the genes within each locus makes accurate assembly and variant detection challenging. These difficulties are particularly challenging in IGK, where large duplicated regions add an additional layer of complexity.

Recent advances in long-read sequencing and haplotype-resolved genome assemblies now provide a powerful platform to revisit these regions. Assemblies such as T2T-CHM13 (6), and a growing number of good quality haplotype-resolved assemblies from individuals of diverse ancestries, offer the resolution needed to systematically explore IGK and IGL gene content and variability.

To ensure consistent and accurate analysis of these assemblies, rigorous and standardized annotation is essential. IMGT®, the international ImMunoGeneTics information system® (http://www.imgt.org) (3), offers a comprehensive framework for the gene classification, functionality, and nomenclature. IMGT resources support consistent annotation and cross-comparison across genomes, offering tools to identify novel genes, alleles, and structural features within complex IG loci.

In this study, we present a comprehensive IMGT analysis of the human IGK and IGL loci, using two high-quality reference assemblies, GRCh38 and T2T-CHM13, along with 142 haploid chromosomal-level assemblies for IGK and 125 for IGL, representing all major human superpopulations: African (AFR), Admixed American (AMR), East Asian (EAS), European (EUR), and South Asian (SAS). This large-scale dataset, representing, to our knowledge, the most extensive collection of assemblies analyzed to date for this purpose, enabled a population-wide investigation of structural variation (SV) in both loci. It allowed us to assess gene presence and prevalence across individuals and ethnicities, while acknowledging potential limitations and the need for even larger-scale studies.

A complete annotation of the T2T-CHM13 and GRCh38 assemblies for the IGL locus, and of the T2T-CHM13 and NA19240_mat_hprc_f2 assemblies for the IGK locus was performed, and revealed previously undescribed IGKV and IGLV genes and alleles, including novel polymorphisms. Many of these new alleles were recurrent across multiple individuals and were supported by read-level evidence and validated using cross-referenced databases.

Our findings expand the IMGT reference directory, provide a refined perspective of the structural variation landscape, showing the prevalence and distribution of SV across a larger and more diverse set of individuals, and offering insights into the diversity and evolutionary dynamics of the human light chain loci. These results provide a valuable resource for future studies of antibody repertoires, population immunogenetics, and immunogenomic diversity in health and diseases, including the role of light chain variation in diagnostics and prognostics, such as in chronic lymphocytic leukaemia and immune responses to infections like COVID-19 and immunological mechanisms underlying cancer (7–9).

## Materials and Methods

The analysis of the new human genomic data was conducted using the IMGT biocuration pipeline, as previously outlined (10). This approach integrates thorough manual curation with a suite of internal software tools, such as IMGT/LIGMotif (11), NtiToVald (12), and IMGT/Automat (12), all based on the IMGT-ONTOLOGY axioms and concepts: “IDENTIFICATION,” “DESCRIPTION,” “CLASSIFICATION,” “NUMEROTATION,” “LOCALIZATION,” “ORIENTATION,” and “OBTENTION”, ensuring accuracy and consistency throughout the annotation process (13).

### Genome Assembly Selection and Filtering

All 554 publicly available human chromosomal assemblies were retrieved from NCBI (14) as of 15 July 2025. The dataset included both reference assemblies (GRCh38 and T2T-CHM13) and additional assemblies from individuals representing diverse populations. Assemblies were initially filtered based on the presence of the chromosome containing the locus of interest; chromosome 2 for the IGK and chromosome 22 for the IGL. NCBI flags such as partial sequence warnings, suspension notices, or diploid assembly status were reviewed, and assemblies were excluded when appropriate.

### Reference Loci IMGT Annotation, Evaluation and Validation

The two high-quality reference assemblies, GRCh38 and T2T-CHM13, were used to generate complete annotations of the IGL locus, and the assemblies T2T-CHM13 and NA19240_mat_hprc_f2 were used for the complete annotation of the the IGK locus.

Corresponding HiFi long-read sequencing data were retrieved from NCBI, when available, and aligned with Minimap2 (15) (v2.24-r1122) for long-read data to the chromosome regions encompassing each locus. Locus quality was evaluated using IMGT/StatAssembly (v1.0.0) (16,17) based on these long-read alignments.

Loci were extracted following the established IMGT biocuration procedure (18) and each was assigned a unique IMGT accession number. Annotation was then performed using IMGT/LIGMotif (11), which provided initial predictions of the genes position and functionality: functional (F), open reading frames (ORFs), and pseudogenes (P) according to the ‘IDENTIFICATION’ axiom of IMGT-ONTOLOGY (19). All gene and allele candidates were then curated manually using Vector NTI (v11.5.2, Thermo Fisher), enabling detailed inspection of gene start and end positions, exon-intron structure, and gene labels in accordance with IMGT criteria.

Novel genes and alleles were identified based on sequence variations from the IMGT reference set and validated through long-read data (for T2T-CHM13 and NA19240_mat_hprc_f2) and/or cross-referencing with external databases such as NCBI whole-genome shotgun contigs (wgs) database using Blast+ (v.2.16.0).

Coding regions were further examined with IMGT/V-QUEST (20) and then the remaining IMGT standardized labels were verified through sequence alignment via BLAST and Multiple Sequence Alignment (MSA) methods, conducted with Clustal Omega tool/EMBL-EBI (21) or MAFFT (22).

All validated sequences, together with updated gene tables and locus representations, were integrated into the IMGT reference directory and web resources (available at: https://www.imgt.org/IMGTrepertoire/, https://www.imgt.org/vquest/refseqh.html and https://www.imgt.org/IMGTgenedbdoc/dataupdates.html)

### IMGT Nomenclature of Novel IGK and IGL Genes and Alleles

The nomenclature of novel IGK and IGL genes and alleles followed the standardized IMGT gene nomenclature, in use since 1988 and detailed in the IMGT-ONTOLOGY “CLASSIFICATION” axiom (https://www.imgt.org/IMGTScientificChart/Nomenclature/IMGTnomenclature.php). This system encodes information about gene type, allele, genomic position and gene duplication status.

New genes and alleles were identified by comparing the coding regions (V, J, or C) of each sequence against the IMGT reference directory. Sequences that corresponded to a known gene at the same relative genomic position (defined by neighboring genes or gene order), but differed by one or more nucleotides, were considered novel alleles according to IMGT criteria. In contrast, sequences with lower identity located between two previously annotated genes were classified as novel genes and named according to their genomic position.

According to IMGT nomenclature standards, IGKV genes in the distal region are named by adding the letter “D” after the subgroup number of their corresponding proximal genes to denote duplicated loci.

### NCBI assemblies from diverse populations: Assembly Curation and Locus Identification

In parallel, all the filtered human chromosome-level haploid assemblies representing diverse human populations (African (AFR), Admixed American (AMR), East Asian (EAS), European (EUR), and South Asian (SAS)) were analysed. As mentioned, all assemblies were downloaded from NCBI and filtered separately for the presence of the assembled chromosome 2 (IGK) and chromosome 22 (IGL). Locus delimitations were determined by comparison to the annotated reference loci and the IMGT reference set. Assemblies that lacked complete locus coverage or exhibited extensive assembly fragmentation in the locus region were excluded from further analysis. The ancestry origins of each assembly and their categorization into superpopulations followed the metadata provided by NCBI and the information supplied by the respective submitters. Notably, the two Saudi Arabian assemblies from one individual, were grouped under EUR due to the absence of a distinct Middle Eastern superpopulation category in our framework.

### Global Assemblies Locus Evaluation and Comparative Gene Analysis

To evaluate the distribution of IGK and IGL genes across assemblies from different ethnicities, a presence/absence analysis was conducted using a custom alignment-based pipeline.

IMGT reference genes retrieved from IMGT/GENE-DB for each locus were first clustered at 98% identity using CD-HIT (v4.8.1) (18) to generate representative sequences and minimize multiple hits for highly similar alleles. These representative sequences were then aligned to the extracted loci using minimap2 (v2.27) (15) with short-read alignment settings (-x sr).

Custom python scripts were used to parse the resulting PAF alignments, apply conservative filtering thresholds (minimum 70% gene coverage and 80% sequence identity), and generate a binary presence/absence matrix. Genes meeting both criteria were marked as present (1); all others were marked as absent (0), resulting in final presence/absence tables for further analysis. For duplicated genes, particularly in the IGK locus, two hits were typically observed: one in the proximal region and one in the distal region, in reverse and forward orientations, respectively. This approach allowed approximate differentiation of duplicated genes and facilitated the construction of the presence/absence matrix.

To validate the results from the short-read (SR) method, BLASTn was performed using longer sequences for each IGK and IGL gene. For this, the V-REGION of each IGK and IGL gene from the T2T assembly was extracted along with 3,000 nucleotides upstream and downstream. This ensured that alignments occurred at the correct genomic positions rather than at highly similar sites, reducing the likelihood of random hits. BLAST searches were conducted between the extracted loci of each studied assembly and these extended T2T reference sequences, using a minimum identity threshold of 98.5%. BLAST results were analyzed using a 60% overlap threshold, retaining only the best match for each position to avoid duplicate results especially for the IGK locus. Exceptions due to closely located genes were manually verified.

All the statistical and visualization analyses were carried out using python and standard scientific libraries.

Assemblies exhibiting gene absences or duplications inconsistent with the reference loci were subjected to additional manual analysis following IMGT biocuration standards. Assemblies that failed verification were excluded from the final dataset, to ensure accurate and reliable gene content across all loci.

## Results

### Curation and Composition of the High-Quality Assembly Dataset for IGK Locus Analysis

As of 15 July 2025, 554 human chromosomal assemblies were available on NCBI. Of these, 362 assemblies were retained for further analysis, primarily due to the presence of chromosome 2, while 126 assemblies lacking chromosome 2 were excluded. Among the retained assemblies, 65 were further removed based on NCBI annotation status, being labeled as partial, suppressed, or diploid (Figure 1A) (Supplementary Table S1, sheet: IGK).

**Figure 1.**
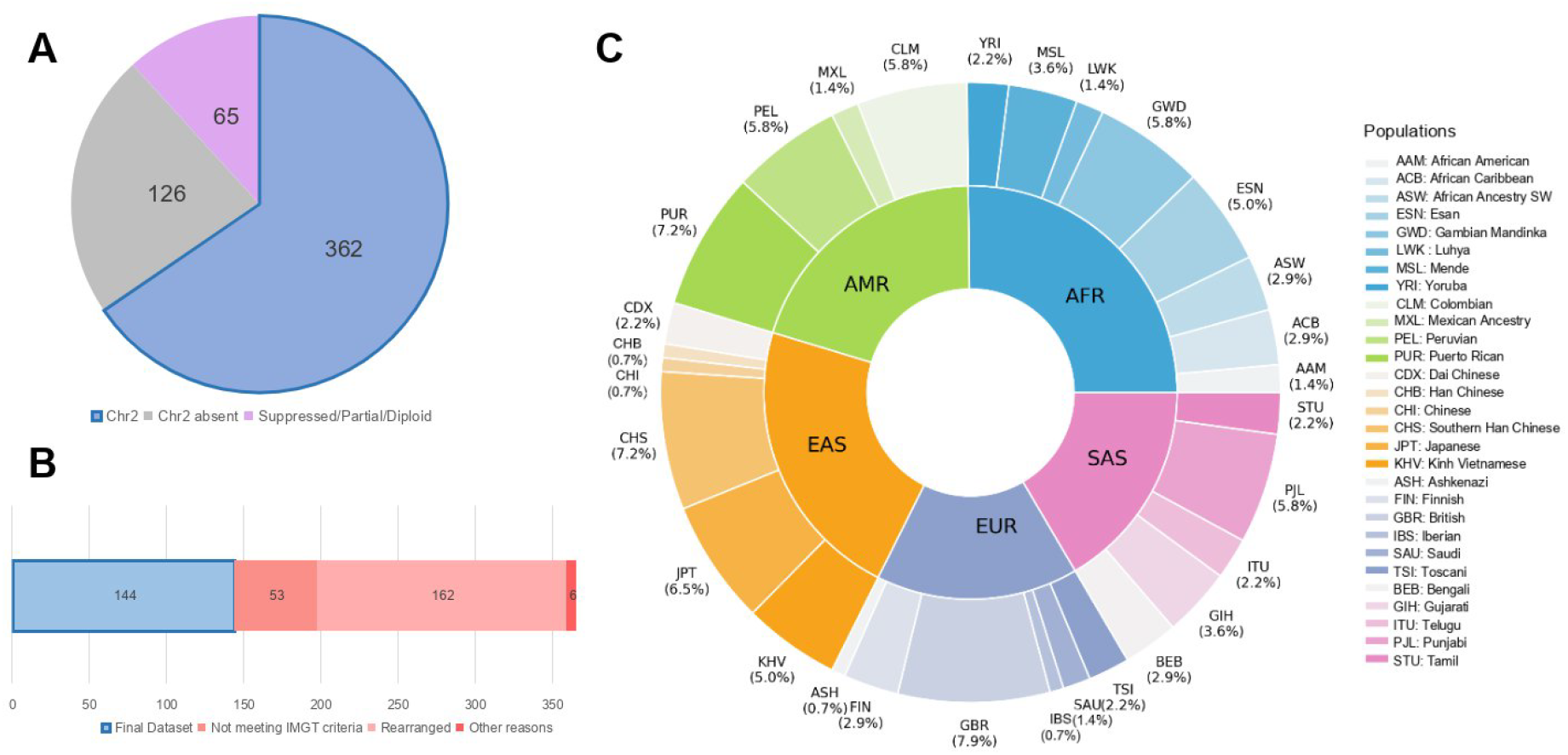
Overview of IGK Dataset Filtering and Population Distribution. **A:** Breakdown of all 554 human chromosomal assemblies in NCBI as of 15 July 2025, categorized by chromosome 2 presence and assembly status. **B:** Stacked bar showing the filtering of the 362 assemblies containing chromosome 2, highlighting the assemblies selected for analysis (144) versus those excluded. **C:** Distribution of the final IGK dataset across superpopulations and populations, as determined from ancestry metadata from NCBI and submitters’ databases.

Among the 362 primarily selected assemblies, 162 were flagged as rearranged or potentially rearranged, particularly when many V and J genes were missing. For the IGK locus, assemblies with rearrangements were excluded from the final dataset, as its inverted duplicated region makes interpretation of rearrangements complex. In addition, 53 assemblies were removed for failing to meet IMGT standards, either due to structural inconsistencies in gene order or gaps affecting locus organization. Unexpectedly, 4 assemblies lacked the IGK locus entirely. In one case (HG03486_pat_hprc_f2), this was due to a partially sequenced chromosome 2, while in the other three (HG04184_pat_hprc_f2, HG02257_mat_hprc_f2, and PGP1v1), the IGK locus was entirely absent. Furthermore, 2 assemblies were removed because they originated from the same individual (HG002.mat.cur.20211005 and HG002.pat.cur.20211005, corresponding to hg002v1.1.mat and hg002v1.1.pat), in these cases, the more recent assemblies were retained. (Figure 1B) (Supplementary Table S1, sheet: IGK).

Following this strict filtering, the final dataset comprises 144 high-quality germline assemblies. Of these, 140 assemblies represent all major human superpopulations, AFR, AMR, EAS, SAS, and EUR, while 2 assemblies lack ancestry metadata (Supplementary Table S1, sheet: IGK) (Figure 1C). The dataset also includes the two reference assemblies (T2T-CHM13v2.0 and GRCh38, https://www.imgt.org/IMGTrepertoire/index.php?section=LocusGenes&repertoire=locus&species=human&group=IGK).

### Comprehensive Annotation of Novel Genes and Alleles in the Human IGK Locus

From the complete annotation of the IGK locus in two assemblies (T2T-CHM13v2.0; IMGT000281 and NA19240_mat_hprc_f2; IMGT000273), 5 novel IGKV genes were identified (3 functional, 2 pseudogenes). Notably, one of the novel functional genes corresponds to the previously non-localized gene IGKV1-NL. This gene was detected between IGKV3D-7 and IGKV1D-8, consistent with prior studies (23), and has been renamed IGKV1D-7-1.

Additionally, the other two novel functional genes, IGKV1-40-1 and its duplicated and inverted IGKV1D-40-1, located upstream of IGKV2-40 and downstream of IGKV2D-40, respectively, confirming previous studies (23). These genes were absent from GRCh38, likely due to sequencing errors, but have now been fully annotated. Finally, 2 novel duplicated genes were identified, IGKV(II)-26-1 in the proximal region (positioned between IGKV2-26 and IGKV1-27) and its duplicate IGKV(II)D-26-1*01 in the distal (positioned between IGKV2D-26 and IGKV1D-27).

Comprehensive analysis of the two IGK loci led to the identification of 10 previously unreported alleles within existing IGKV and IGKJ genes (IMGT/GENE-DB program version 3.1.42, 2024-09-25). These comprise 9 IGKV (4 F, 5 P), and 1 IGKJ (1 ORF) alleles.

All novel genes and alleles have been fully annotated in the sequences IMGT000225 and IMGT000280, and are considered high-confidence, supported by sequencing read data and have been found in multiple individuals in the NCBI wgs datasets.

### Structural and Gene Variation at the IGK Locus Across Populations and Ancestral Backgrounds

Across the 144 assemblies analyzed, the IGK locus showed a highly conserved gene content and organization (Supplementary Table S2, sheet: IGK, Figure 2A). Despite this overall conservation, the novel gene IGKV1D-7-1 was identified as a structural variant, as it was detected in only 30 assemblies and absent from the remaining 114 (Figure 2A). As shown in the stacked bar plot of Figure 2B, the frequencies varied across superpopulations, with the gene present in 5.5% of EUR assemblies (1/22), 12.5% of SAS (3/24), and 10.7% of AMR (3/28), whereas higher prevalence was observed in AFR and EAS assemblies, with 31.4% (11/35) and 35.5% (11/31) of assemblies carrying the gene, respectively.

**Figure 2.**
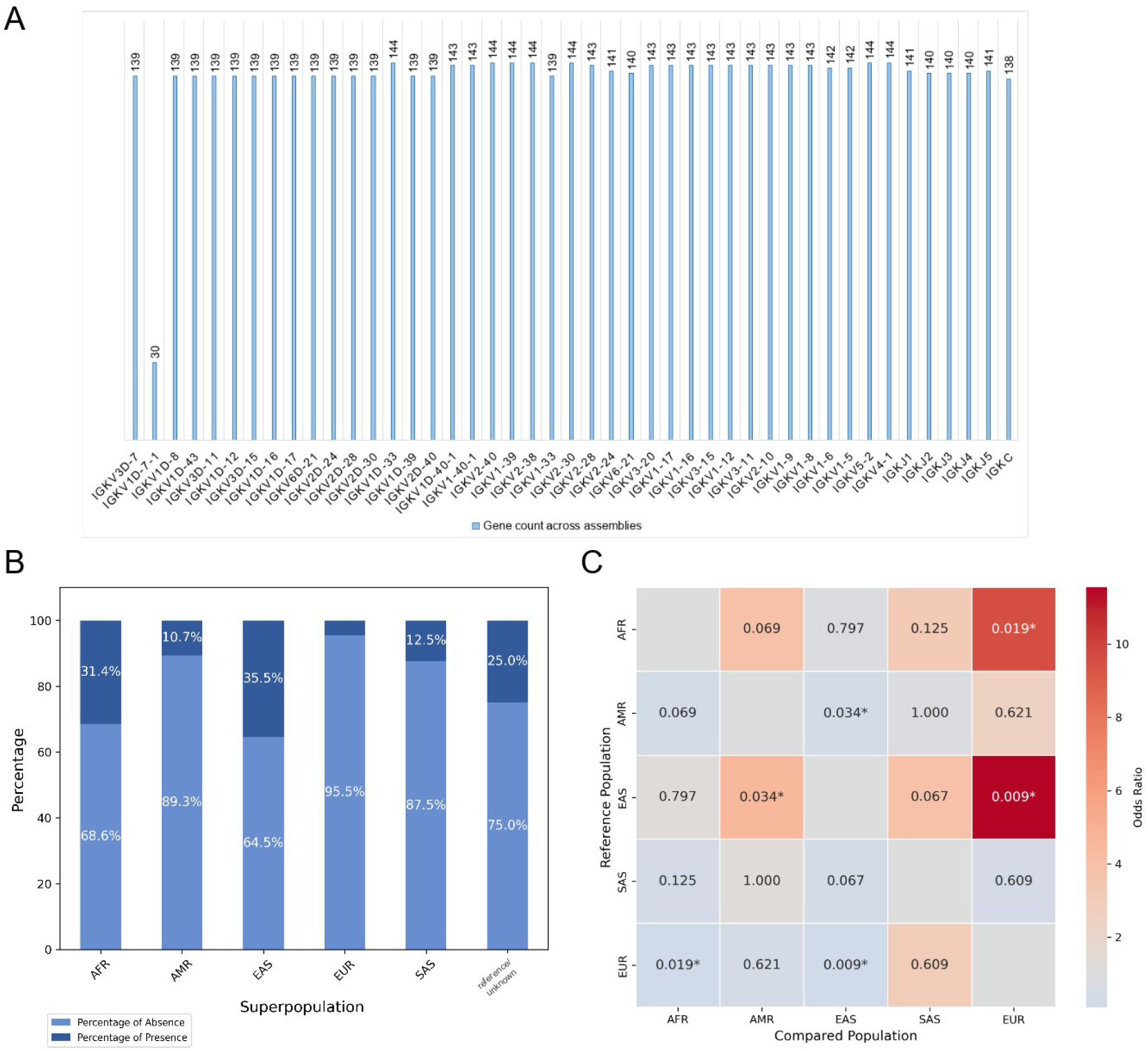
IGKV1D-7-1 Gene Presence and Population Differences. **A:** The graph illustrates the frequency of each functional IGKV gene across assemblies. **B:** Stacked bar plot showing the proportion of assemblies carrying the IGKV1D-7-1 gene across human superpopulations. **C:** Heatmap of pairwise odds ratios for gene presence between populations. Odds ratios >1 indicate a higher likelihood of gene presence in the reference population relative to the comparison population. Statistically significant differences (Fisher’ s exact test, p < 0.05) are marked with asterisks, and the corresponding p-values are shown within each cell.

To assess whether these differences were population-specific, pairwise Fisher’s exact tests were performed, and odds ratios and p-values were calculated. The results were visualized in a heatmap (Figure 2C), with each cell showing the corresponding p-value. Odds ratios greater than 1 indicated a higher likelihood of gene presence in the reference population compared with the comparison population, and statistically significant associations (p < 0.05) were marked with an asterisk.

The analysis showed that AFR and EAS assemblies were significantly more likely to carry IGKV1D-7-1 than EUR, consistent with the previously observed prevalence. EAS assemblies also had a higher likelihood of carrying the gene compared with AMR. In contrast, comparisons between AFR and AMR and EAS and SAS did not reach statistical significance (p ≈ 0.07), which may reflect the limited sample sizes.

A rare structural variation involving the complete deletion of the distal region of the IGK locus was observed in 5 out of 144 assemblies, including 2 EUR, 1 AMR, 1 EAS, and 1 SAS assembly. This deletion results in the absence of multiple distal IGKV genes, potentially reducing the diversity of the IGK repertoire in these individuals.

Additional regions with gene deletions were observed, each detected in only one assembly (Supplementary Table S2, sheet: IGK). For example, the longest deletion was identified in CM087187.1, where genes from IGKV2-29 to IGKV1-5 were not detected by any method.

Other notable deletions were observed in approximately the same region, including IGKV1-27 to IGKV6-21 in CM087317.1, IGKV2-26 to IGKV6-21 in CM086975.1, and IGKV2-23 to IGKV6-21 in CM089428.1. No assembly gaps were identified in these regions, and the corresponding genes remained undetected by all applied methods, suggesting either true structural variation or potential sequencing artifacts.

### Curated Dataset of High-Quality Assemblies of Different Ancestries for IGL Locus Annotation and Novel Gene Discovery

Of the 554 chromosomal assemblies available on NCBI as of 15 July 2025, 412 assemblies were excluded; 361 due to absence of chromosome 22 and 60 due to NCBI annotation status as partial, suppressed, or diploid (Figure 3A). For the remaining 133 haploid chromosomal assemblies, we processed to IGL locus localization, with only 41 identified within the expected chromosomal range based on reference sequence (21-24Mb on chromosome 22, https://www.imgt.org/IMGTrepertoire/LocusGenes/locusdesc/human/IGL/Hu_IGLdesc.html) and 1 assembly lacked the locus entirely (Figure 3B). Of the final 132 assemblies containing the IGL locus, one paternal assembly (CM085807.1, HG03834) was excluded for incomplete locus representation, and four others (ex. CH003517.1) due to disrupted gene order and quality inconsistent with IMGT criteria (Supplementary Table S1, sheet: IGL). The final 127 high-quality assemblies represented all major human superpopulations, including 29 AFR, 17 AMR, 35 EAS, 7 SAS, and 27 EUR individuals, while 12 assemblies were missing ancestry metadata (Supplementary Table S2, sheet: IGL) (Figure 3C). Six of them contained rearranged loci (three EUR, two EAS, and one AFR), originated from B lymphocyte cell lines, and were retained due to their relevance in capturing biologically meaningful stages of somatic VJ recombination (24).

**Figure 3.**
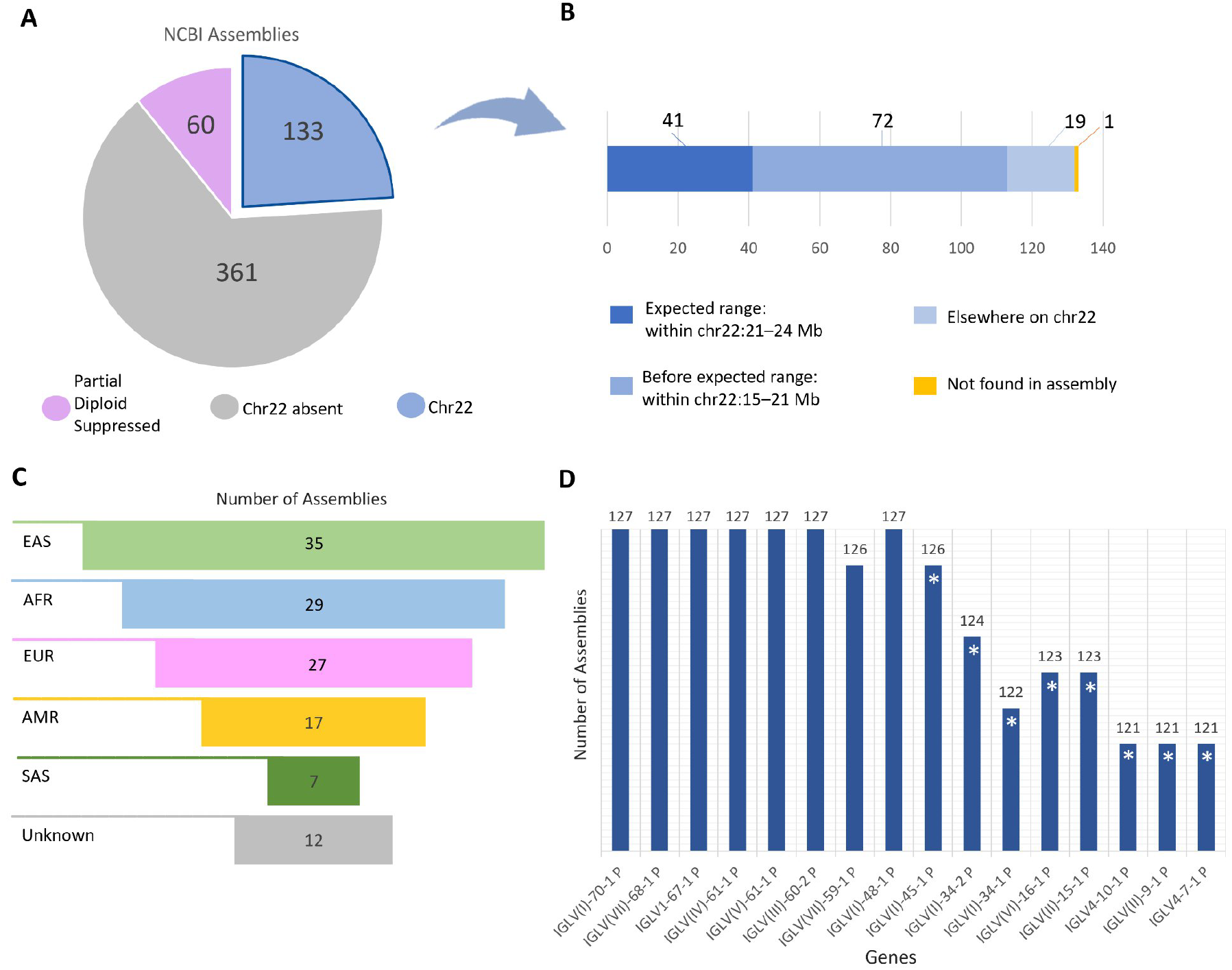
Curation of high-quality assemblies and identification of novel IGL genes across diverse ancestries. **A:** Distribution of all 554 human chromosomal assemblies available in NCBI as of 15 July 2025, according to chromosome 22 availability and assembly status. **B:** Localization of the IGL locus among 133 assemblies containing chromosome 22. The expected locus range (21-24Mb) is defined based on GRCh38 and T2T-CHM13 references. Assemblies are classified according to locus position or complete absence. Detailed information on the locus location of each assembly can be found on Supplementary Table S2, sheet: IGL. **C:** Superpopulation distribution of the final 127 assemblies retained for IGL analysis, assigned using ancestry metadata from NCBI and submitters’ databases. **D:** Frequency of the 16 novel IGLV genes identified across the 127 assemblies. Genes are ordered by genomic position within the locus. An asterisk indicates absence in a minority of cases, likely due to somatic rearrangement.

Sixteen novel IGLV genes were identified by annotation of both the two reference assemblies and the 125 additional high-quality haploid assemblies from diverse superpopulations (IMGT000225 and IMGT000280). In addition, comprehensive annotation of the two reference loci revealed 22 new alleles within existing IGLV genes (IMGT/GENE-DB program version:3.1.42 (2024-09-25)). The novel genes were consistently identified across the dataset, with the majority present in nearly all assemblies (Figure 3D). Their absence in a small number of cases, as indicated with an asterisk in Figure 3D, is attributable to somatic rearrangement, which obscures the original germline configuration in those assemblies (Supplementary Table S2, sheet: IGL). All 16 new genes are annotated in the 2 reference sequences with corresponding alleles (IMGT000225 and IMGT000280) and are supported by presence in one or more individuals in the NCBI nucleotide collection (nr/nt) or NCBI wgs.

### Gene Variations and Gene Presence/Absence Patterns Across the IGL Locus

The overall gene content and organization of the IGL locus were highly conserved across the 127 assemblies, with most deviations representing individual cases rather than systematic population-specific patterns. However, one subgroup 5 variable gene, IGLV5-39, exhibited recurrent absence in 75 out of 127 assemblies (Supplementary Table S2, sheet: IGL). Although IGLV5-39 is a F gene, it was missing in a substantial number of assemblies. Its presence was confirmed in 52 assemblies, where it was consistently localized between IGLV(I)-42 and IGLV(VII)-41-1. Figure 4A summarizes the presence–absence distribution of IGLV5-39, revealing a pronounced and consistent gene loss, while also illustrating individual and superpopulation-level variation. The graph suggests that EUR and AMR genomes have roughly equal likelihood of carrying the gene, while the other populations are more likely to lack it. Pairwise Fisher’s exact tests (Figure 4B) did not reach statistical significance (p > 0.05), indicating that, while differences in IGLV5-39 presence were observed between populations, these differences could not be confirmed with the current sample sizes.

**Figure 4.**
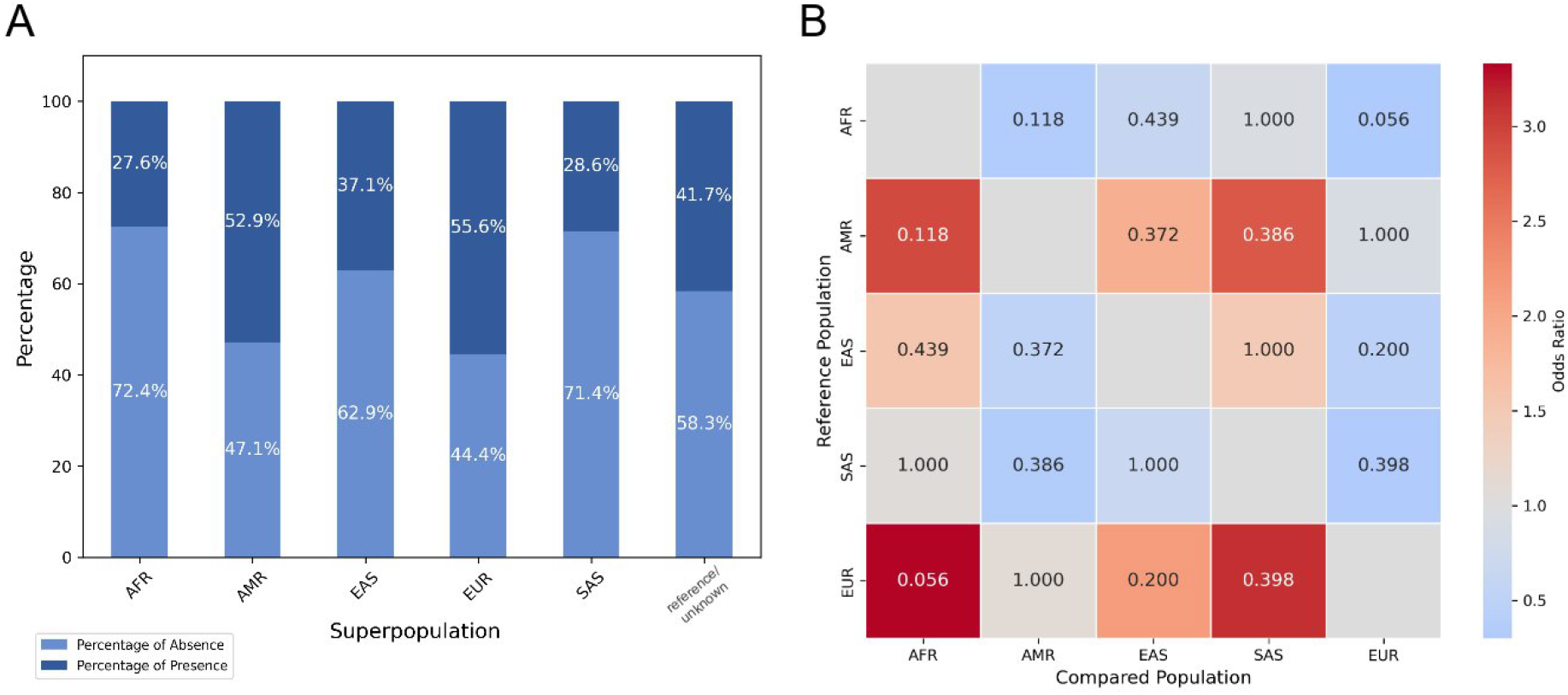
IGLV5-39 Gene Presence and Population Variations. **A:** Stacked bar showing the proportion of assemblies carrying the IGLV5-39 gene across human superpopulations. **C:** Heatmap of pairwise odds ratios for gene presence between populations. Odds ratios >1 indicate a higher likelihood of gene presence in the reference population relative to the comparison population, with corresponding p-values displayed within each cell.

Additional polymorphisms were detected in specific individuals. For instance, the paternal assembly of individual HG00544 (Southern Han Chinese, CM089381.1) lacked 11 consecutive IGLV3 and IGLV2 subgroup genes and clans (IGLV3-21 to IGLV2-14), with no sequence gaps in the region (∼9.3 kb), suggesting a possible structural deletion, while the maternal assembly of the individual did not show this deletion. A similar case was found in NA20850 (Gujarati Indian, CM094315.1), where a block of seven genes was absent between IGLV4-60 and IGLV11-55, spanning 13 kb, compared to 52 kb in the reference genome.

Interestingly, examination of the constant region confirmed potential copy number variations (CNV) (25), with multiple assemblies exhibiting duplications of J–C clusters (e.g., J2–C2 or J3–C3), as well as triplications involving J2–C2 alone (e.g., CM092025.1). Ambiguity in distinguishing IGLC2 and IGLC3 arises from their high sequence similarity (99%), making it difficult to determine whether these represent IGLC2 duplications or novel C3 variants. For example, in CM101478.1 (NA18970, unknown ancestry, geographic origin Japan) and CM101699.1 (NA18943, Japanese), three copies of IGLC2*02 were identified with no IGLC3 present, potentially indicating a IGLC3 variant identical in sequence to IGLC2. Other assemblies (e.g., CM009893.1, Southern Han Chinese) exhibited duplication of entire J2–C2/J3–C3 blocks, with junction and constant regions showing near-identical sequences. These structural configurations are documented in Supplementary Table S2, sheet: IGL, with selected cases underscored to highlight representative examples. Manual verification of a subset of assemblies confirmed the presence of duplicated or triplicated constant genes, supporting the occurrence of potential CNVs at the IGL constant region.

### Comparison of Gene Detection by Short-Read Mapping and BLASTn

To assess the reliability of gene detection across assemblies, two complementary approaches were employed. Minimap2 (v2.27) with short-read alignment settings (-x sr) successfully detected the majority of immunoglobulin genes across assemblies. BLASTn, using longer reference sequences, was then applied to validate these results and confirm the genes identified by sr mapping. In addition, a few genes that were missed due to the strict thresholds in SR mapping were identified by BLAST and are indicated with asterisks in Supplementary Table S2. For example, in the IGK locus, IGKV3-7 in assembly CM086905.1 was not detected by SR mapping but was captured by BLAST. Similarly, while IGKJ1, IGKJ2, IGKJ4, and IGKJ5 were consistently detected by both methods, IGKJ3 was identified only by BLAST. Overall, this combined approach highlights the complementary strengths of SR mapping for high-confidence detection and BLAST for increased sensitivity in challenging regions.

## Discussion

The comprehensive analysis of both IGK and IGL loci across multiple human assemblies revealed a complex landscape of genetic variation, including novel genes, allelic polymorphisms, and structural variants. Importantly, the use of IMGT positional nomenclature was critical to confirm the existence of genes and to distinguish different alleles, ensuring consistency and accuracy in gene annotation. Examination of these loci across diverse populations highlighted differences in gene presence and frequency, providing insights into the population-specific architecture of light chain immunoglobulin repertoires. This study represents, to our knowledge, the most extensive dataset to date investigating structural variation and population-level differences in the IGK (n=144) and IGL (n=127) loci of human. This comprehensive analysis provides valuable insights into population-specific patterns of structural variations, revealing differences in gene presence and frequencies across diverse human genome assemblies. Further data will be essential to capture rare variants and comprehensively characterize population-specific differences, ultimately providing a more complete picture of immunogenetic variation.

In this study, to assess gene presence and absence in each genome, we employed two complementary approaches: short-read mapping using minimap2 and sequence alignment with BLAST using longer reference sequences. Our analysis demonstrated that SR mapping is highly effective not only for read alignment but also for mapping genes directly on assembled genomes, as the majority of genes detected by SR were confirmed with BLAST. Occasional discrepancies between SR mapping and BLAST highlight the strengths and limitations of each approach. SR mapping provides rapid and high-confidence detection for most genes, but strict thresholds can exclude certain sequences, particularly in highly similar and short genes such as the J genes. BLAST, by aligning longer reference sequences, increases sensitivity and allows recovery of genes missed by SR mapping. These findings underscore the complementary nature of combining short-sequence and long-sequence alignment methods for a comprehensive characterization of immunoglobulin loci.

### Key Insights into IGK Locus: Variation, Comparative Insights across Populations and Their Immunogenetic Implications

In this study, we performed a comprehensive analysis of the human IGK locus across 144 haploid assemblies from diverse populations, including GRCh38 and T2T-CHM13, to assess gene content, allelic diversity, and structural variation.

Previous studies have suggested that the IGK locus is largely complete and accurately annotated (26). However, in our analysis of 2 haploid assemblies using the IMGT biocuration pipeline, we identified 5 previously unreported IGKV genes and 10 novel alleles. Together with recent reports of additional IGK polymorphism (23,27), these findings indicate that the IGK locus may still harbor undiscovered variation, particularly when including genomes from diverse ancestral backgrounds.

Among the novel alleles identified in our study, IGKV2D-40*02 is of particular interest, as it has been implicated in diverse pathological processes. IGKV2D-40 has been reported as a candidate biomarker in tendon healing and venous thrombosis (28), and has also been linked to prognosis in head and neck squamous cell carcinoma (29) and colorectal cancer (30), with potential as an immunotherapy target. While the functional consequences of the IGKV2D-40*02 polymorphism remains to be determined, variation in this gene could potentially influence protein function, immune regulation, and disease outcomes.

Similarly, we identified a novel SNP in IGKV2D-28 (IGKV2D-28*02), that has been previously reported as a potential biomarker for gestational diabetes mellitus (GDM) (31). While its precise effects remain unknown, this SNP may influence gene activity and expression.

Together with IGKV2D-40*02, these findings highlight the relevance of novel IGKV alleles in immunogenetic variation and disease pathways, underscoring the importance of further research across broader cohorts.

In addition to single-nucleotide variation, structural variation also contributes to IGK diversity. An SV insertion in the distal region corresponding to IGKV1D-7-1 was detected in approximately 30 assemblies, confirming previous reports (23)). Our results highlight the variable presence of IGKV1D-7-1 across populations, with higher frequencies observed in AFR and EAS assemblies compared with EUR, SAS, and AMR, suggesting that population-specific structural variation contributes to IGK diversity. Pairwise comparisons confirmed that AFR and EAS genomes are significantly more likely to carry IGKV1D-7-1 relative to EUR, and EAS more likely than AMR. Other comparisons, including SAS versus AFR or EAS, did not reach statistical significance, even though the stacked bar plot indicated relatively low frequencies of IGKV1D-7-1 in SAS. These non-significant results may reflect the limited sample sizes in these populations. Overall, these findings illustrate how structural variation at the IGK locus can differ across populations and highlight the value of large, diverse datasets for uncovering population-specific patterns.

In addition to this insertion, we observed a rare SV involving the complete deletion of the distal region of the IGK locus in 5 out of 144 assemblies, that has been previously reported (32). This deletion results in the loss of multiple distal IGKV genes, potentially reducing the diversity of the IGK repertoire in affected individuals.

Finally, several putative structural variant (SV) deletions were detected in individual assemblies, potentially representing rare SVs within the IGK locus. Of particular interest were deletions spanning the region between IGKV1-27 and IGKV6-21, in which one assembly lacked all 8 intervening genes, another lacked 6, and a third lacked 3. The repeated observation of deletions within this region may indicate the presence of a potential genomic hotspot. Further validation in larger datasets is required to determine whether these events represent rare structural variants or sequencing artifacts.

### Key Insights into IGL Locus: Variation, Comparative Insights across Populations and Their Immunogenetic Implications

Our study provides an updated and comprehensive characterization of the human IGL locus using 127 haploid assemblies from diverse populations, including assemblies GRCh38 and T2T-CHM13. Through the IMGT biocuration pipeline, we identified 16 new IGLV genes and 22 novel alleles, extending the known IGL germline repertoire and illustrating ongoing gene and allele discovery and refinement.

The IGL locus typically spans between 21 and 24 Mb on chromosome 22 and it lies within a structurally complex region enriched in segmental duplications and low-copy repeats (33). As a result, the IGL locus appeared shifted, fragmented, or absent in several of the haploid assemblies, likely reflecting both genuine structural variation and assembly challenges. Despite this, a substantial number of complete loci allowed detailed annotation and confirmation of gene content.

Among the new entries are IGLV(I)-34-1 and IGLV(I)-34-2, located within a previously described 11.9 kb intergenic region between IGLV7-35 and IGLV2-34 (34). We confirm this insertion spans ∼12.9 kb and harbors two P genes consistently present in most assemblies including the reference genomes. The formerly unlocalized IGLV2-NL1, now designated IGLV2-34, is now fully annotated and observed in all unrearranged loci (34,35).

Several previously ambiguous or under-characterized IGL genes were clarified in this study, with IGLJ6, previously suspected to be P, identified in nearly all assemblies and consistently annotated as F in the IMGT databases (36). Similarly, IGLV3-12, IGLV3-22, and IGLV4-3, which appeared at lower frequencies in transcriptomic studies, were consistently present at the genomic level, supporting their inclusion in the germline repertoire (36). Conversely, IGLV5-39 demonstrated limited representation across the dataset, consistent with prior reports of polymorphic insertions, deletions, or highly divergent segments in subgroup 5 (37–39). In our dataset, we confirm the presence of all IGLV subgroups, including less-characterized such as IGLV9 (IGLV9-49) and IGLV10 (e.g., IGLV10-64) (40) and the consistent presence of clinically relevant genes including IGLV1-44, IGLV6-57, and IGLV2-23. Notably, IGLV6-57, associated with AL amyloidosis, was found in 129 assemblies, supporting its consistent germline presence and highlighting the need for further study of allelic diversity and regulatory features (40–42).

In line with previous findings, we observed that the IGL locus displays relatively low allelic diversity, especially compared to the very complex IGH, indicating that light chain diversity is under tighter evolutionary constraint to reduce the risk of self-reactivity (38,43). Indeed, genes previously described as monomorphic, such as those in the IGLV7 subgroup showed minimal allelic diversity (except IGLV7-46, with 5 alleles).

Emerging evidence indicates that structural variation in the IGLJ and IGLC genes is more prevalent than previously appreciated (41,43–45). In our study, duplications of IGLC2 and IGLC3 were detected across diverse ancestries, consistent with earlier findings in Japanese, Tunisian, Lebanese and African, likely reflecting historical recombination events with potential expression-level effects (46).

The refined IMGT IGL reference set has important implications for immunogenetics and clinical research, as such variability contributes to antibody diversity and disease susceptibility to both malignant and infectious diseases. Specific IGLV gene usage has been associated with prognosis in B cell malignancies, such as chronic lymphocytic leukemia and AL amyloidosis, as well as immune responses to infections, including SARS-CoV-2 (9,40,47).

Overall, our analyses of the IGK and IGL loci reveal previously unrecognized allelic and structural variation, emphasizing the value of diverse population datasets to capture the full spectrum of immunogenetic diversity. Our results underscore the impact of new germline variants in enhancing repertoire analysis and providing a framework to investigate the genetic basis of IG variation in cancer, autoimmunity, and emerging infections, thereby advancing our understanding of antibody diversity, immune regulation and vaccine responsiveness. Future studies integrating high-quality diverse assemblies and functional data will be critical to fully understanding the contribution of light chain diversity to human health.

## Data Availability

IMGT^®^ is freely available online for academics and non-profit use at http://www.imgt.org/. All the databases and tools referred to in this article are accessible from the IMGT^®^ website. The data underlying this article are available in the article and in its online supplementary materials.

## Supplementary Data

Supplementary Table S1 ‘NCBI Assemblies Information’

Supplementary Table S2: ‘S1.Genes presence-absence matrix’

Supplementary Figure S1 ‘IGK and IGL holistic locus representation’

## Acknowledgments

We sincerely appreciate the IMGT^®^ team for their unwavering dedication and enthusiasm. Over the course of this approximately, several Erasmus+ and French university internship students contributed, most for a duration of two months. We value the energy and fresh perspectives they brought to the team and we list them in alphabetical order: Arthur Appelgren, Flore Brantschen, Theodora Giannousa, Eve Gun.

## Author Contributions

Conceptualization, Methodology, Validation, and writing – review & editing: M.G., A.P., G.F., J.J.-M., G.Z., P.D., V.G., and S.K.; Investigation, Data Curation, Visualization and Writing – original draft: A.P. and M.G.; Supervision: V.G. and S.K. Funding acquisition: S.K. All authors have read and agreed to the published version of the manuscript.

## References

1. Lefranc MP, Lefranc G. IMGT®Homo sapiens IG and TR Loci, Gene Order, CNV and Haplotypes: New Concepts as a Paradigm for Jawed Vertebrates Genome Assemblies. Biomolecules. 2022 Mar;12(3):381.

2. Lefranc MP. Nomenclature of the human immunoglobulin lambda (IGL) genes. Exp Clin Immunogenet. 2001;18(4):242–54.

3. Manso T, Folch G, Giudicelli V, Jabado-Michaloud J, Kushwaha A, Nguefack Ngoune V, et al. IMGT® databases, related tools and web resources through three main axes of research and development. Nucleic Acids Res. 2022 Jan 7;50(D1):D1262–72.

4. Kim DY, To R, Kandalaft H, Ding W, van Faassen H, Luo Y, et al. Antibody light chain variable domains and their biophysically improved versions for human immunotherapy. mAbs. 2014;6(1):219–35.

5. Schneider VA, Graves-Lindsay T, Howe K, Bouk N, Chen HC, Kitts PA, et al. Evaluation of GRCh38 and de novo haploid genome assemblies demonstrates the enduring quality of the reference assembly. Genome Res. 2017 May;27(5):849–64.

6. Nurk S, Koren S, Rhie A, Rautiainen M, Bzikadze AV, Mikheenko A, et al. The complete sequence of a human genome. Science. 2022 Apr;376(6588):44–53.

7. Tobin G. The Immunoglobulin genes and Chronic Lymphocytic Leukemia (CLL). Ups J Med Sci. 2005;110(2):97–114.

8. Gao H, Yu L, Yan F, Zheng Y, Huang H, Zhuang X, et al. Landscape of B Cell Receptor Repertoires in COVID-19 Patients Revealed Through CDR3 Sequencing of Immunoglobulin Heavy and Light Chains. Immunol Invest. 2022 Oct 3;51(7):1994–2008.

9. Gudowska-Sawczuk M, Moniuszko-Malinowska A, Pączek S, Guziejko K, Chorąży M, Mroczko B. Evaluation of Free Light Chains (FLCs) Synthesis in Response to Exposure to SARS-CoV-2. Int J Mol Sci. 2022 Sept 30;23(19):11589.

10. Pégorier P, Bertignac M, Chentli I, Nguefack Ngoune V, Folch G, Jabado-Michaloud J, et al. IMGT® Biocuration and Comparative Study of the T Cell Receptor Beta Locus of Veterinary Species Based on Homo sapiens TRB. Front Immunol. 2020;11:821.

11. Lane J, Duroux P, Lefranc MP. From IMGT-ONTOLOGY to IMGT/LIGMotif: the IMGT standardized approach for immunoglobulin and T cell receptor gene identification and description in large genomic sequences. BMC Bioinformatics. 2010 Apr 30;11:223.

12. Folch G, Jabado-Michaloud J, Bellahcene F, Regnier L, Giudicelli V, Lefranc MP. IMGT/Automat: the strategy for the annotation of human and mouse cDNA nucleotide sequences of IG and TR. Nat Preced. 2009 Apr 23;1–1.

13. Giudicelli V, Lefranc MP. IMGT-ONTOLOGY 2012. Front Genet. 2012;3:79.

14. Sayers EW, Bolton EE, Brister JR, Canese K, Chan J, Comeau DC, et al. Database resources of the national center for biotechnology information. Nucleic Acids Res. 2022 Jan 7;50(D1):D20–6.

15. Li H. New strategies to improve minimap2 alignment accuracy. Bioinformatics. 2021 Oct 8;37(23):4572–4.

16. Zeitoun G, Debbagh C, Georga M, Papadaki A, Sideri I, Folch G, et al. IMGT/StatAssembly [Internet]. Zenodo; 2025 [cited 2025 July 29]. Available from: https://zenodo.org/records/15396812

17. Gaoussou Sanou, Guilhem Zeitoun, Taciana Manso, Milad Eidi, Shamsa Batool, Anjana Kushwaha, et al. IMGT ® at scale: FAIR, Dynamic and Automated Tools for Immune Locus Analysis. in press;

18. Pégorier P, Bertignac M, Chentli I, Nguefack Ngoune V, Folch G, Jabado-Michaloud J, et al. IMGT® Biocuration and Comparative Study of the T Cell Receptor Beta Locus of Veterinary Species Based on Homo sapiens TRB. Front Immunol [Internet]. 2020 May 5 [cited 2025 July 29];11. Available from: https://www.frontiersin.org/journals/immunology/articles/10.3389/fimmu.2020.00821/full

19. Frontiers | IMGT-ONTOLOGY 2012 [Internet]. [cited 2025 Sept 22]. Available from: https://www.frontiersin.org/journals/genetics/articles/10.3389/fgene.2012.00079/full

20. Brochet X, Lefranc MP, Giudicelli V. IMGT/V-QUEST: the highly customized and integrated system for IG and TR standardized V-J and V-D-J sequence analysis. Nucleic Acids Res. 2008 July 1;36(Web Server issue):W503–508.

21. Madeira F, Park Y mi, Lee J, Buso N, Gur T, Madhusoodanan N, et al. The EMBL-EBI search and sequence analysis tools APIs in 2019. Nucleic Acids Res. 2019 July 2;47(W1):W636–41.

22. Katoh K, Rozewicki J, Yamada KD. MAFFT online service: multiple sequence alignment, interactive sequence choice and visualization. Brief Bioinform. 2019 July 19;20(4):1160–6.

23. Engelbrecht E, Rodriguez OL, Shields K, Schultze S, Tieri D, Jana U, et al. Resolving haplotype variation and complex genetic architecture in the human immunoglobulin kappa chain locus in individuals of diverse ancestry. Genes Immun. 2024 Aug;25(4):297–306.

24. ten Boekel E, Melchers F, Rolink A. The status of Ig loci rearrangements in single cells from different stages of B cell development. Int Immunol. 1995 June;7(6):1013–9.

25. Taub RA, Hollis GF, Hieter PA, Korsmeyer S, Waldmann TA, Leder P. Variable amplification of immunoglobulin lambda light-chain genes in human populations. Nature. 1983 July 14;304(5922):172–4.

26. Collins AM, Wang Y, Singh V, Yu P, Jackson KJ, Sewell WA. The reported germline repertoire of human immunoglobulin kappa chain genes is relatively complete and accurate. Immunogenetics. 2008 Nov 1;60(11):669–76.

27. Khatri I, Berkowska MA, van den Akker EB, Teodosio C, Reinders MJT, van Dongen JJM. Population matched (pm) germline allelic variants of immunoglobulin (IG) loci: Relevance in infectious diseases and vaccination studies in human populations. Genes Immun. 2021;22(3):172–86.

28. Saarensilta A, Chen J, Reitzner SM, Hart DA, Ahmed AS, Ackermann PW. Novel tissue biomarker candidates to predict both deep venous thrombosis and healing outcome after Achilles tendon rupture. Sci Rep. 2025 Mar 1;15(1):7318.

29. Huang C, Liu J. Identification of the Immune Cell Infiltration Landscape in Head and Neck Squamous Cell Carcinoma (HNSC) for the Exploration of Immunotherapy and Prognosis. Genet Res. 2022 Dec 28;2022:6880760.

30. Lin K, Huang J, Luo H, Luo C, Zhu X, Bu F, et al. Development of a prognostic index and screening of potential biomarkers based on immunogenomic landscape analysis of colorectal cancer. Aging. 2020 Mar 31;12(7):5832–57.

31. Payot MD, Villavieja A, Pineda-Cortel MR. Preliminary Investigation of Potential Early Biomarkers for Gestational Diabetes Mellitus: Insights from PTRPG and IGKV2D-28 Expression Analysis. Int J Mol Sci. 2024 Sept 30;25(19):10527.

32. Pargent W, Schäble KF, Zachau HG. Polymorphisms and haplotypes in the human immunoglobulin kappa locus. Eur J Immunol. 1991 Aug;21(8):1829–35.

33. Demaerel W, Mostovoy Y, Yilmaz F, Vervoort L, Pastor S, Hestand MS, et al. The 22q11 low copy repeats are characterized by unprecedented size and structural variability. Genome Res. 2019 Sept;29(9):1389–401.

34. Watson CT, Steinberg KM, Graves TA, Warren RL, Malig M, Schein J, et al. Sequencing of the human IG light chain loci from a hydatidiform mole BAC library reveals locus-specific signatures of genetic diversity. Genes Immun. 2015;16(1):24–34.

35. Williams SC, Winter G. Cloning and sequencing of human immunoglobulin V lambda gene segments. Eur J Immunol. 1993 July;23(7):1456–61.

36. Mikocziova I, Peres A, Gidoni M, Greiff V, Yaari G, Sollid LM. Germline polymorphisms and alternative splicing of human immunoglobulin light chain genes. iScience. 2021 Oct 22;24(10):103192.

37. Lefranc MP. The immunoglobulin factsbook. San Diego: Academic Press; 2001. xiv+457. (Factsbook series).

38. Lefranc MP, Pallarès N, Frippiat JP. Allelic polymorphisms and RFLP in the human immunoglobulin lambda light chain locus. Hum Genet. 1999 May;104(5):361–9.

39. Moraes Junta C, Passos GAS. Genomic EcoRI polymorphism and cosmid sequencing reveal an insertion/deletion and a new IGLV5 allele in the human immunoglobulin lambda variable locus (22q11.2/IGLV). Immunogenetics. 2003 Apr;55(1):10–5.

40. Solomon A, Weiss DT. Structural and functional properties of human lambda-light-chain variable-region subgroups. Clin Diagn Lab Immunol. 1995 July;2(4):387–94.

41. Gibson WS, Rodriguez OL, Shields K, Silver CA, Dorgham A, Emery M, et al. Characterization of the immunoglobulin lambda chain locus from diverse populations reveals extensive genetic variation. Genes Immun. 2023 Feb;24(1):21–31.

42. Ozaki S, Abe M, Wolfenbarger D, Weiss DT, Solomon A. Preferential Expression of Human λ-Light-Chain Variable-Region Subgroups in Multiple Myeloma, AL Amyloidosis, and Waldenström’s Macroglobulinemia. Clin Immunol Immunopathol. 1994 May 1;71(2):183–9.

43. Kay PH, Moriuchi J, Ma PJ, Saueracker E. An unusual allelic form of the immunoglobulin lambda constant region genes in the Japanese. Immunogenetics. 1992 Mar 1;35(5):341–3.

44. van der Burg M, Barendregt BH, van Gastel-Mol EJ, Tümkaya T, Langerak AW, van Dongen JJM. Unraveling of the Polymorphic Cλ2-Cλ3 Amplification and the Ke+Oz™ Polymorphism in the Human Igλ Locus. J Immunol. 2002 July 1;169(1):271–6.

45. Blasini A, Delgado M, Valdivieso C, Guevara P, Ramirez J, Stekman I, et al. Review: Restriction fragment length polymorphisms of constant region genes of immunoglobulin lambda chains in Venezuelan patients with systemic lupus erythematosus. Lupus. 1996 Aug 1;5(4):300–2.

46. Ghanem N, Dariavach P, Bensmana M, Chibani J, Lefranc G, Lefranc MP. Polymorphism of immunoglobulin lambda constant region genes in populations from France, Lebanon and Tunisia. Exp Clin Immunogenet. 1988;5(4):186–95.

47. Shao Z, Feng Y, Zhong L, Xie Q, Lei M, Liu Z, et al. Clinical efficacy of intravenous immunoglobulin therapy in critical ill patients with COVID-19: a multicenter retrospective cohort study. Clin Transl Immunol. 2020;9(10):e1192.

