## Supplementary Figure S1 for "Novel Genes and Polymorphisms in Human Immunoglobulin Light Chains Across Diverse Populations Through Comprehensive IMGT Analysis"

**A.**

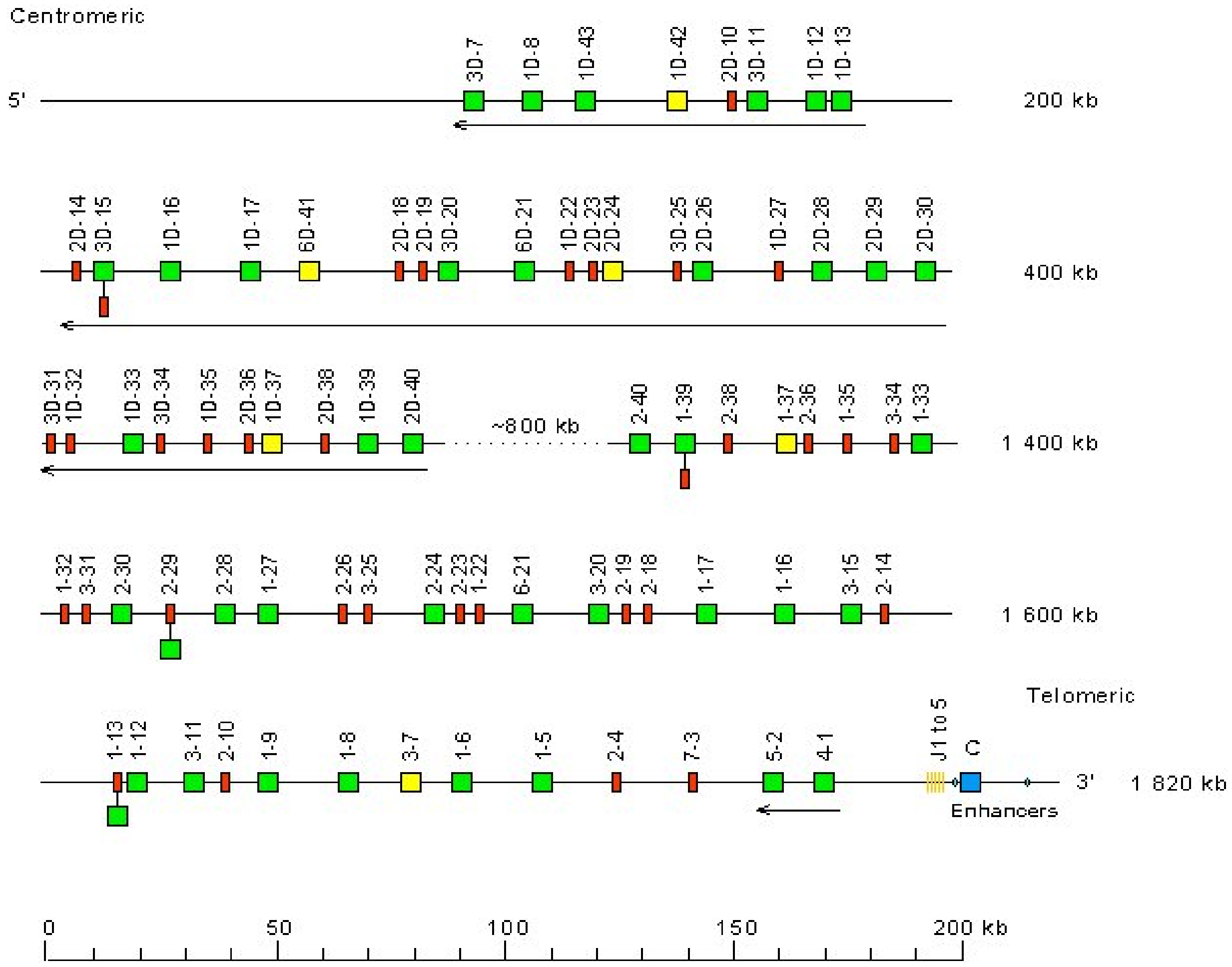

Centromeric
5'
3D-7
1D-8
1D-43
1D-42
2D-10
3D-11
1D-12
1D-13
200 kb
2D-14
3D-15
1D-16
1D-17
6D-41
2D-18
2D-19
3D-20
6D-21
1D-22
2D-23
2D-24
3D-25
2D-26
1D-27
2D-28
2D-29
2D-30
400 kb
3D-31
1D-32
1D-33
3D-34
1D-35
2D-36
1D-37
2D-38
1D-39
2D-40
~800 kb
2-40
1-39
2-38
1-37
2-36
1-35
3-34
1-33
1 400 kb
1-32
3-31
2-30
2-29
2-28
1-27
2-26
3-25
2-24
2-23
1-22
6-21
3-20
2-19
2-18
1-17
1-16
3-15
2-14
1 600 kb
Telomeric
1-13
1-12
3-11
2-10
1-9
1-8
3-7
1-6
1-5
2-4
7-3
5-2
4-1
J1 to 5
C
3'
1 820 kb
Enhancers
0
50
100
150
200 kb

**B.**

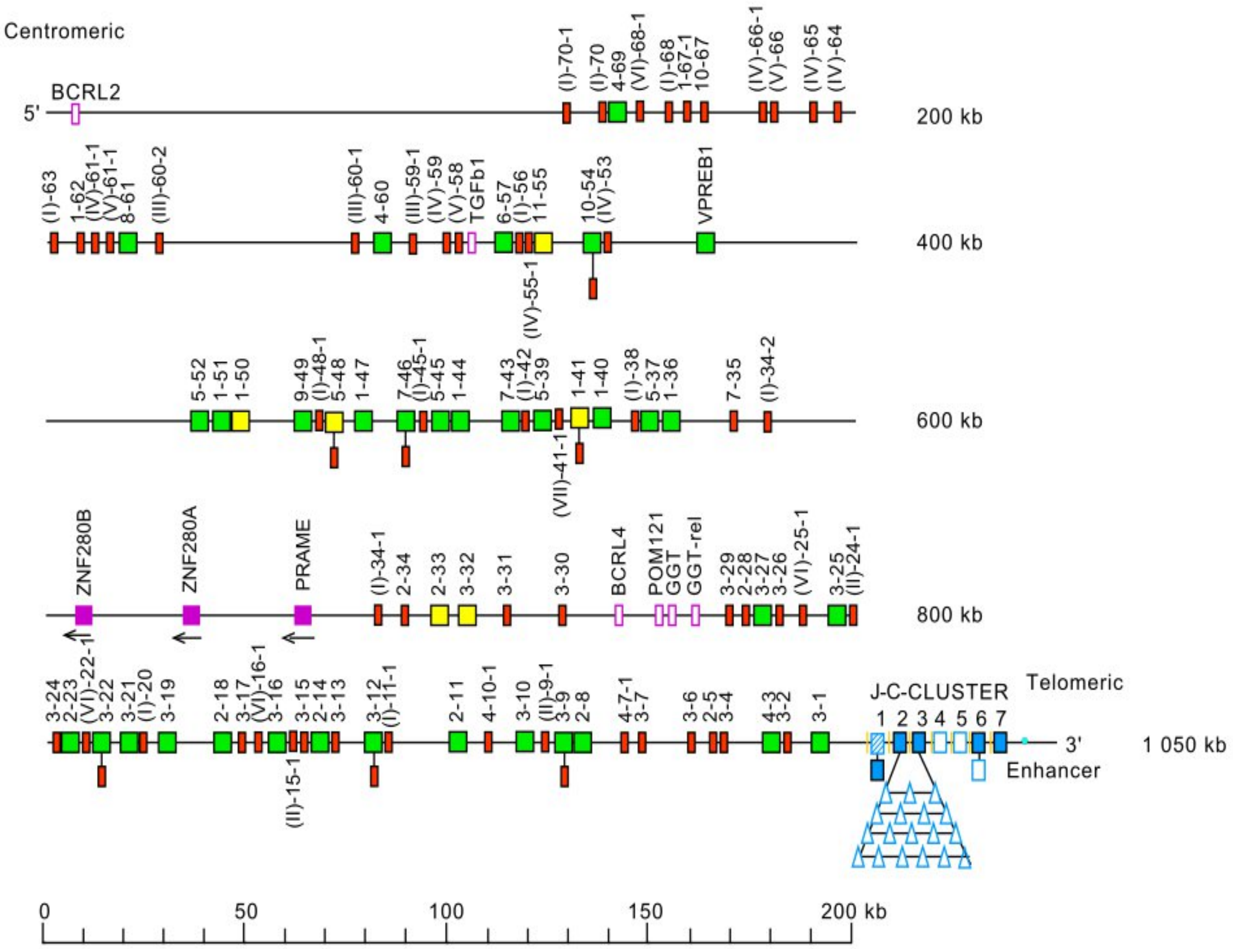


**Supplementary Figure S1.** Holistic IMGT® schematic representation of the human IGK (**A**) and IGL (**B**) loci. This figure provides a comprehensive overview of the organization of the human light chain loci reflecting the functional and structural complexity of the locus. IGK locus, located on chromosome 2 (2p11.2), contains IGKV, IGKJ and IGKC genes, while IGL locus, on chromosome 22 (22q11.2), comprises IGLV, IGLJ, and IGLC genes. Each gene is depicted as a box, with its color indicating functionality and gene type, consistent with the IMGT® color menu (https://www.imgt.org/IMGTScientificChart/RepresentationRules/colormenu.php#LOCUS). Genes shown with two boxes indicate cases where the gene has been identified with different functionalities in various alleles. This representation is continuously updated on the IMGT® webpage (https://www.imgt.org/IMGTrepertoire/LocusGenes/locusdesc/human/IGL/Hu_IGLdesc.html) as new genes or new alleles functionalities are identified. Arrows indicate IGKV genes oriented opposite to the J–C cluster.
